# Meta-Analysis Identifies Pleiotropic Loci Controlling Phenotypic Trade-offs in Sorghum

**DOI:** 10.1101/2020.10.27.355495

**Authors:** Ravi V. Mural, Marcin Grzybowski, Chenyong Miao, Alyssa Damke, Sirjan Sapkota, Richard E. Boyles, Maria G. Salas Fernandez, Patrick S. Schnable, Brandi Sigmon, Stephen Kresovich, James C. Schnable

**Affiliations:** Center for Plant Science Innovation and Department of Agronomy and Horticulture, University of Nebraska-Lincoln, Lincoln, NE USA; Department of Plant Pathology, University of Nebraska-Lincoln, Lincoln, NE USA; Advanced Plant Technology Program, Clemson University, Clemson, SC USA; Department of Plant and Environment Sciences, Clemson University, Clemson, SC USA; Pee Dee Research and Education Center, Clemson University, Florence, SC, USA; Department of Agronomy, Iowa State University, Ames, IA USA; Feed the Future Innovation Lab for Crop Improvement Cornell University, Ithaca, NY USA

## Abstract

Community association populations are composed of phenotypically and genetically diverse accessions. Once these populations are genotyped, the resulting marker data can be reused by different groups investigating the genetic basis of different traits. Because the same genotypes are observed and scored for a wide range of traits in different environments, these populations represent a unique resource to investigate both pleiotropy and genotype by environment interactions. Here we assembled a set of 234 separate trait datasets for the Sorghum Association Panel, a group of 406 sorghum genotypes widely employed by the sorghum genetics community. Comparison of genome wide association studies conducted with two independently generated marker sets for this population demonstrate that existing genetic marker sets do not saturate the genome and likely capture only 35-43% of potentially detectable loci controlling variation for traits scored in this population. While limited evidence for pleiotropy was apparent in cross-GWAS comparisons, a multivariate adaptive shrinkage approach recovered both known pleiotropic effects of existing loci and new pleiotropic effects, particularly significant impacts of known dwarfing genes on root architecture. In addition, we identified new loci with pleiotropic effects consistent with known trade-offs in sorghum development. These results demonstrate the potential for mining existing trait datasets from widely used community association populations to enable new discoveries from existing trait datasets as new, denser genetic marker datasets are generated for existing community association populations.

## Introduction

The value of common mapping populations in diverse species has been recognized by quantitative genetics for decades. These common populations can be genotyped by a single research group, and genetically identical individuals distributed to the larger research community which could score traits of interest and test for associations between published genetic markers and trait values across the population. Early populations included sets of recombinant inbred lines (RILs)^1^. RIL populations were developed, genotyped and released to the maize^2, 3^ and arabidopsis^4^ genetics communities. Mapping quantitative trait loci (QTL) in RIL populations provided relatively high power to detect variants with even small numbers of markers as large scale linkage disequilibrium (LD) permits the identification of associations between genetic markers and trait values for markers at quite some distance from the causal variant. Yet the high LD of RIL populations also meant that researchers were unlikely to identify associations with the causal gene or variant without generating new follow-up populations for fine mapping. Improvements to genotyping technologies ultimately permitted the use of natural populations with more rapid decay of LD^5–7^. Like earlier RIL populations, association populations can be distributed among research groups, permitting association mapping and later genome-wide association studies (GWAS) to be conducted for multiple traits without the need to generate new marker data.

The number of markers needed to saturate the genome of a target population for GWAS is determined by the size of the species’ genome, the speed at which LD decays within the target population, and the minimum level of LD between a causal variant and a genotyped marker where statistically significant associations will still be detected, which in turn depends on the proportion of total population variance explained by the causal variant^8^. An early estimate based on 1,122 genetic markers suggests that >100,000 genetic markers would be necessary to conduct GWAS in populations spanning global sorghum diversity when target causal variants explained >10% of total trait variance and >350,000 genetic markers when target causal variants explained 5-10% of total trait variance^8^. Estimates of this type are quite sensitive to the rate of LD decay. Estimates of the average distance at which LD decays below an *r^2^* value of 0.1 in sorghum range from 10 kilobases to 350 kilobases depending on the population and genetic marker set employed^8–13^. LD also varies among different portions of the genome, creating an addition challenge to accurate simulation.

Higher marker densities increase the odds of identifying causal variants that explain more modest proportions of the total variance for a target trait. When new sets of genetic marker become available for existing association populations it is possible to reanalyze previously collected trait datasets. In addition, studies based on simulated data suggest multivariate analyses may increase true positive rates relative to trait-by-trait univariate GWAS^14^. Multivariate analysis may provide value in case where individual causal loci have pleiotropic effects on multiple traits which are measured separately. The degree of pleiotropy for loci influencing variation in quantitative traits in plants remains uncertain. A study of maize leaf traits found little incidence of pleiotropy^15^ while detecting modest evidence of pleiotropy between the elongation of leaves tassel and ears^16^. The sorghum QTL Atlas, a meta-analysis of reported QTL map locations from 146 publications of diverse RIL populations identified QTL hotspots on chromosome 2, corresponding to the *brown nucellar layer2* (*B2*) gene in sorghum controlling, and chromosome 7, corresponding to *dwarf3* (dw3)^17^. However, the relatively large confidence interval of QTL peaks identified via mapping in RIL populations can make it difficult to determine whether QTL hotspots represent a single highly pleiotropic gene or multiple linked genes^18^. In maize, the pleiotropic effects of a large effect QTL for plant architecture on ear traits was long thought to be explained by a single gene, *teosinte branched1* (*tb1*), but was later fractionated into multiple partially linked loci for ear related traits^19^.

The Sorghum Association Panel (SAP) was first assembled in 2008^20^. After some additions, the population ultimately consisted of 40621 lines selected to represent the global genetic diversity of sorghum. Because it was intended that this population be grown and phenotyped in the temperate United States, the majority of the lines included in the panel were generated by the Sorghum Conversion program^20,22^. Sorghum from many parts of the world fail to flower during the summer growing season in the temperate United States. Sorghum Conversion lines are the result of crossing diverse sorghum germplasm from around the world to a temperate adapted donor parent (BTx406) and then recurrently backcrossing the progeny to the exotic-tropical parent for four generations while selecting for temperate adaptation, including flowering during the summer in temperate latitudes and short stature^23^. Retrospective genomic analysis of many Sorghum Conversion lines identified three genomic intervals where the haplotype of the donor parent is over represented in the population. These three intervals corresponded to the locations of dwarfing genes *dw1-dw3*^24^. No comparable independent intervals were identified for loci conferring photoperiod insensitivity, however *maturity1* (*ma1*) has a large selection sweep on Chr6 that is linked to *dwarf2* (dw2)^24^.

Initially the SAP was genotyped for only a set of 49 SSR markers^20^. Subsequently, the SAP was employed for a number of genetic association tests using increasing numbers of markers^20,25–28^. In 2013, a set of several hundred thousand genetic markers was generated for the population using conventional genotyping by sequencing^29,30^. From 2013 onward the SAP was widely employed for GWAS by a range of research groups targeting different traits (Table 1). Here we employ a set of both published and previously unpublished trait datasets from the SAP and multiple genetic marker datasets^30,31^ to empirically evaluate both the degree of saturation achieved by current genetic marker sets and the degree to which detectable loci controlling phenotypic variation in the SAP tend to be pleiotropic or non-pleiotropic using a multi-trait approach based on meta-analysis and adaptive shrinkage^32^.

**Table 1.** Papers Scoring Traits in the Sorghum Association Population

| Reference | Year Published | Study Type | Phenotypes Scored | # of SAP Accessions Evaluated | Genetic Associations? | Trait Data Online? |
| --- | --- | --- | --- | --- | --- | --- |
| Casa et al.<sup>20</sup> | 2008 | Vegetative | 8 | 377 | AS | No |
| Brown et al.<sup>25</sup> | 2008 | Height & Inflorescence | 6 | 378 | AS | Yes |
| Vandenbrink et al.<sup>33</sup> | 2010 | Biomass Composition | 2 | 377 | No | Yes |
| Mutava et al.<sup>34</sup> | 2011 | Drought Stress | 10 | 300 | No | No |
| Sukumaran et al.<sup>26</sup> | 2012 | Grain Quality | 10 | 300 | AS | No |
| Wu et al.<sup>27</sup> | 2012 | Tannins | 1 | 161 | AS | Yes |
| Morris et al. A<sup>11</sup> | 2013 | Flavonoids | 2 | 259-387 | GWAS | Yes |
| Morris et al. B<sup>30</sup> | 2013 | Various | 6 | 355 | GWAS | Used public data |
| Hufnagel et al.<sup>28</sup> | 2014 | Phosphorous Deficiency | 10 | 287 | AS | No |
| Kong et al.<sup>35</sup> | 2014 | Tillering and inflorescence | 9 | 377 | GWAS | No |
| Perez et al.<sup>36</sup> | 2014 | Plant Architecture | 6 | 315 | GWAS | Released Here |
| Rhodes et al.<sup>37</sup> | 2014 | Polyphenols | 3 | 308 | GWAS | No |
| Adeyanju et al.<sup>38</sup> | 2015 | Disease Resistance | 6 | 300 | GWAS | Yes, but IDs ambiguous |
| Lasky et al.<sup>39</sup> | 2015 | Drought Stress | 28 | 267 | GWAS | Yes |
| Prom et al.<sup>40</sup> | 2015 | Disease Resistance | 3 | 177 | No | Yes |
| Queiroz et al.<sup>41</sup> | 2015 | Grain Quality | 12 | 100 | No | Yes |
| Li et al.<sup>42</sup> | 2015 | Height | 3 | 307 | GWAS | Used public data |
| Zhang et al.<sup>43</sup> | 2015 | Height & Inflorescence | 12 | 354 | GWAS | No |
| Zhang et al.<sup>44</sup> | 2015 | Seed Size | 6 | 354 | GWAS | Yes |
| Boyles et al.<sup>45</sup> | 2016 | Various | 13 | 378 | GWAS | Released Here |
| Shakoor et al.<sup>46</sup> | 2016 | Elemental Abundance | 22 | 407 | GWAS | Yes |
| Zhao et al.<sup>47</sup> | 2016 | Plant Architecture | 9 | 315 | GWAS | Released Here |
| Boyles et al.<sup>48</sup> | 2017 | Grain Quality | 10 | 378 | GWAS | Released Here |
| Chen et al.<sup>49</sup> | 2017 | Heat Stress | 2 | 374 | GWAS | No |
| Chopra et al.<sup>50</sup> | 2017 | Heat and Cold Stress | 12 | 300 | GWAS | Yes |
| Salas Fernandez et al.<sup>51</sup> | 2017 | Vegetative (HTP) | 4 | 307 | GWAS | No |
| Ortiz et al.<sup>52</sup> | 2017 | Photosynthesis/Cold Stress | 24 | 304 | GWAS | No |
| Paiva et al.<sup>53</sup> | 2017 | Elemental Abundance | 18 | 100 | No | Yes |
| Rhodes et al.<sup>54</sup> | 2017 | Polyphenols | 3 | 266 | GWAS | Yes |
| Rhodes et al.<sup>55</sup> | 2017 | Grain Quality | 4 | 265 | GWAS | Yes |
| Cuevas et al.<sup>56</sup> | 2018 | Disease Resistance | 2 | 335 | GWAS | Yes |
| Breitzman et al.<sup>57</sup> | 2019 | Vegetative (HTP) | 6 | 325 | GWAS | Released Here |
| Cuevas et al.<sup>58</sup> | 2019 | Disease Resistance | 2 | 331 | GWAS | Yes |
| McMaster et al.<sup>59</sup> | 2019 | Mycotoxin | 2 | 98 | No | Yes, but IDs ambiguous |
| Moghimi et al.<sup>60</sup> | 2019 | Cold Stress | 13 | 351 | GWAS | Yes |
| Olatoye et al.<sup>61</sup> | 2019 | Various | 4 | 334 | GWAS | Used public data |
| Prom et al.<sup>62</sup> | 2019 | Disease Resistance | 1 | 359 | GWAS | Yes |
| Zheng et al.<sup>63</sup> | 2019 | Root Architecture (HTP) | 12 | 294 | GWAS | Yes |
| Zhou et al.<sup>64</sup> | 2019 | Inflorescence (HTP) | 8 | 302 | GWAS | Yes |
| Miao et al.<sup>65</sup> | 2020 | Height | 1 | 357 | GWAS | Yes |

## Results

### Properties of SAP trait datasets

A literature review identified 40 papers in which phenotypic data were collected and published from the Sorghum Association Panel (SAP) – or subsets of this population. Of these 40 papers, it was possible to obtain trait values for individual lines in 25 cases. These included 20 papers for which data were provided as supplemental information and 5 papers for which the data was obtained directly from the authors (Table 1). These 25 papers reported data for a total of 190 traits, although it should be noted that some of these traits are similar or identical measurements conducted in different environments or years. Twelve additional phenotypes were derived based on sums or ratios of trait means (Table S1). Previously unpublished data for 19 traits collected in Nebraska and 13 previously unpublished trait datasets collected in South Carolina were also added to the dataset (Table S1). Thus, the final dataset consisting of 234 distinct sets of trait data scored for all or subsets of a common sorghum population. Values of all 234 sets of trait data, including those previously unpublished are provided as supplementary data with this paper. A number of studies reported data from plants grown in growth chambers or other controlled environment conditions. However, the majority of published trait datasets came from field trials in six states – Georgia, Iowa, Kansas, Nebraska, South Carolina and Texas – within the United States with additional trait data collected in Brazil (Figure 1A).

**Figure 1.**
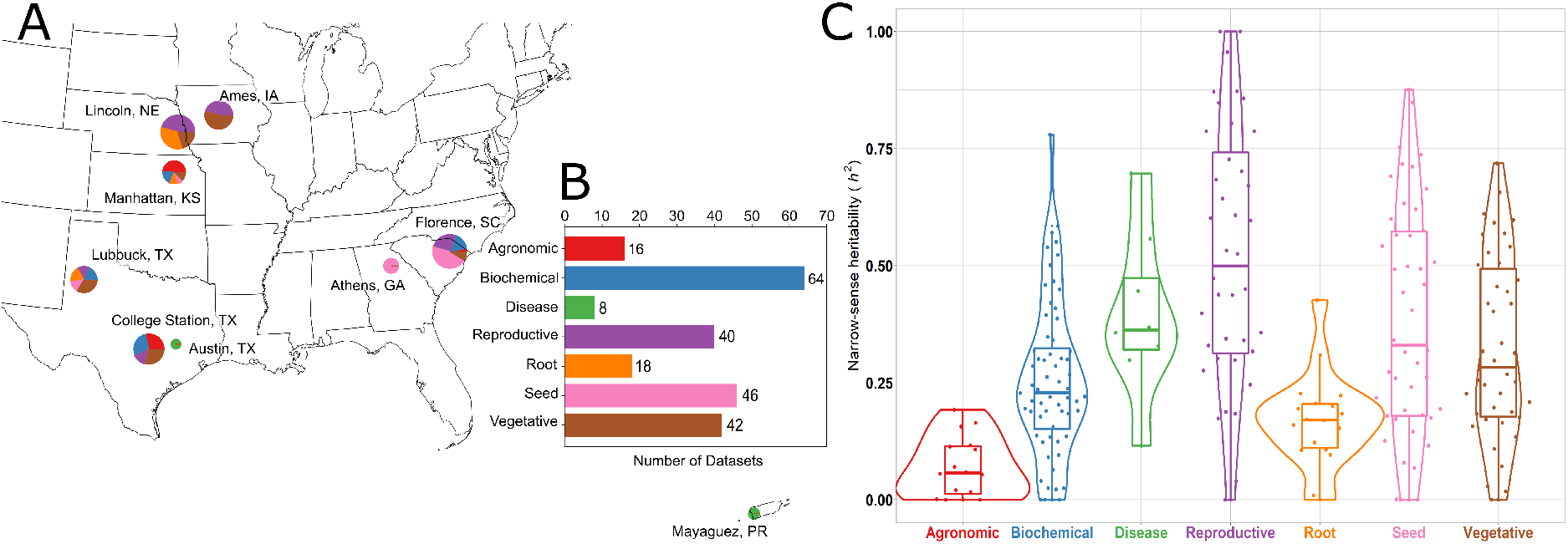
Characteristics of Sorghum Association Panel trait datasets. A) Geographic distribution of trials where trait datasets were collected. Size of circles indicates number of traits collected at a specific geographic location. Colors of circles indicate types of trait datasets collected at that location. Labels for which colors correspond to which types of traits are given in Panel B. A set of 30 traits scored in Nova Porteirinha, Minas Gerais, Brazil^41,53^ are not visible in this panel. B) Representation of seven broad phenotypic categories among the 234 traits collected here. Category assignments for individual traits are provided in Table S1. C) Distributions of narrow sense heritability values, calculated using the 2020 genetic marker dataset^31^, across the same seven broad phenotypic categories are shown in panel B.

Traits could be broadly classified into seven categories including agronomic phenotypes, biochemical phenotypes, disease related phenotypes, root phenotypes, above ground vegetative phenotypes (of which 15 plant height or plant height proxies), reproductive phenotypes, and seed phenotypes (Figure 1B; Table S1). Most studies provided only trait means or best linear unbiased predictor (BLUP)^66^ values (Table S1). As a result, it was not possible to estimate broad sense heritabilities for most traits. However, it was possible to estimate narrow sense heritabilities. Estimates of narrow sense heritability had a median of 0.265 (Figure 1C). Traits with high narrow sense heritability tended to be those related to panicle morphology, grain composition and disease, while traits with estimates of narrow sense heritability close to zero tended to be collected from seedlings, a subset of biochemical traits, and measures of some plasticity across environments (Table S1). Trait datasets belonging to the same categories, as well as trait datasets collected as part of the same studies tended to be correlated more with each other than with other pairs of traits (Figure S1).

### Current sorghum marker sets do not achieve saturation for linkage to causal loci

Previous estimates of the number of markers required to achieve saturation of a sorghum GWAS population have been largely based on simulation studies^8^. The existence of two distinct genetic marker datasets provides an opportunity to empirically estimate the number of markers required to saturate the sorghum genome. In 2013 a set of 265,487 markers on version 1 of the sorghum genome were generated using conventional genotyping by sequencing for 971 lines including 355 members of the SAP^29,30^ (referred to as the “2013 marker set” below). In 2020 a set of 569,305 markers on version 3 of the sorghum genome were generated using a modified genotyping by sequencing strategy for 343 members of the SAP^31,67^ (referred to as the “2020 marker set” below). A set of 304 lines were included in both the 2013 and the 2020 genetic marker datasets. Different protocols were employed to generate these two datasets and these two protocols sequence different subsets of the sorghum genome. As a result, the sets of specific genetic markers genotyped in each dataset should be largely non-overlapping. If current marker datasets are sufficient to saturate the sorghum genome, using different marker datasets would be expected to identify signals from the same regions of the genome. However, if current marker sets are insufficient to saturate the sorghum genome, using different marker datasets to analyze the same trait datasets would be expected to identify only partially overlapping sets of genomic intervals for the same trait datasets.

Each genetic marker dataset was employed to conduct GWAS for each of the 234 trait datasets. For each genetic marker dataset, imputation and filtering based on minor allele frequency specifically within the set of 304 shared lines were conducted using a common set of criteria (See Methods). Similarly, a common trait data processing protocol was employed for each of the 234 trait datasets. This protocol incorporated normalization and outlier removal. GWAS for two of the 234 trait datasets produced questionable results, including distributions of observed p-values inconsistent with expectations, when analyzed with one of the marker sets (Figure S2A-D). The GWAS results for these two trait datasets were removed from downstream analyses for both marker sets.

Among the remaining 232 trait datasets, 36 trait datasets produced at least one significant marker-trait associations (MTAs) when analyzed using the 2013 marker dataset (N=48 significant peaks). These 48 total significant peaks in individual GWAS with 2013 dataset localized to 26 unique regions of the genome as a result of repeated identification of the same genomic intervals in analyses of different trait datasets (Figure 2A). When the same 232 trait datasets were analyzed using the 2020 genetic marker dataset, 40 trait datasets produced at least one significant peak (N=52 significant peaks). These 52 total significant peaks localized to 32 unique locations on the sorghum genome (Figure 2B). In analyses with each of the two genetic marker datasets, multiple distinct signals were observed in the same region of chromosome six including the canonical signal for *dw2*, as well as a separate peak for nickel abundance (both genetic marker datasets), seed weight (2013 genetic marker dataset) and stem size (2020 genetic marker dataset). The clustering of separate peaks may reflect increased statistical power resulting from elevated minor allele frequencies in this interval from over representation of the BTx406 haplotype among sorghum conversion lines in this region^24^.

**Figure 2.**
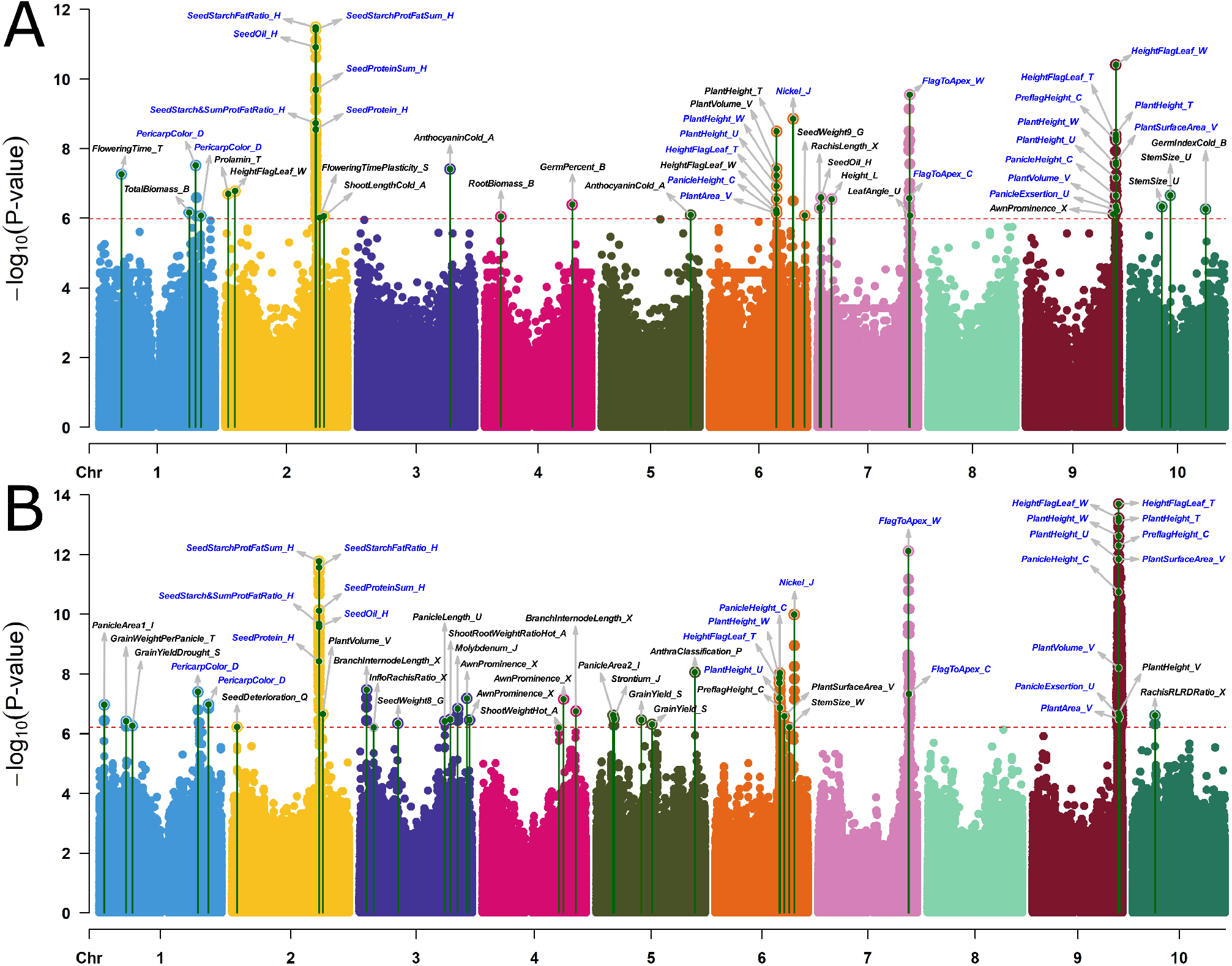
Combined Manhattan plots comparing MTAs identified using different marker datasets for the same individuals: A) Combined Manhattan plot for 36 traits with at least one significant GWAS hit when analyzed using the 2013 genetic marker dataset and considering data from only those 304 sorghum lines genotyped in both the 2013 and 2020 datasets. Green lines topped with circles indicate the physical position and −log10 p-value for the single most significant SNP within a GWAS peak identified for a particular trait. Text labels for individual traits employ trait names provided in Table S1. Dashed red line indicates the cutoff for statistical significance calculated from the effective SNP number in the 2013 genetic marker dataset. B) Combined Manhattan plot for 40 traits with at least one significant GWAS hit when analyzed using the 2020 genetic marker dataset and considering data from only those 304 sorghum lines genotyped in both the 2013 and 2020 datasets. Locations and p-values of the most significant SNP within each peak and statistical significance cut-off labeled as above. Blue labels indicate peaks shared between datasets.

A total of 54 traits exhibited at least one significant trait associated marker, with 36 traits exhibiting at least one peak in the 2013 dataset and 40 traits exhibiting at least one peak in the 2020 dataset. Among these 54 traits a total of 22 traits exhibited at least one significant peak that was shared between the two marker datasets, while there were 14 unique traits in the 2013 marker set and 18 unique traits in the 2020 marker data set, each of which exhibited at least one unique peak (Figure S3A-B). Of the 22 traits which exhibited at least one significant peak when analyzed with each of the genetic marker datasets, nine were height traits. Among the 32 traits which exhibited at least one significant peak when analyzed with one genetic marker dataset and no significant peaks when analyzed with the other, only two were height related traits. The non-representative nature of height related traits may be explained both by the presence of three segregating large effect mutations for height in this population and the large LD blocks which exist around these genes as a result of selection during the temperate adaptation process^24^. Excluding height related trait datasets, 13 trait datasets produced at least one significant MTA, when analyzed with either genetic marker dataset and 30 trait datasets produce at least one significant MTA, when analyzed with one and only one of the genetic marker datasets (Figure S3C).

However, even when at least one significant MTA was identified when the same trait dataset was analyzed with each of the two genetic marker datasets, these MTAs may not correspond to the same causal loci. Of 74 unique MTAs between a given trait dataset and a given genomic interval identified using the two genetic marker datasets, 26 of the same MTAs were identified using both genetic marker datasets, 22 were identified only using the 2013 dataset and 26 were identified using only the 2020 dataset (Figure 2; S3D-E). Peaks associated with plant height were disproportionately likely to be identified in analyses using both genetic marker datasets. Nineteen total peaks associated with plant height traits were identified of which 13 were identified when the same trait was analyzed with either genetic marker dataset. Excluding plant height-related traits, 55 distinct MTAs were identified between variation in a trait dataset and a given region of the genome. A total of 13 MTAs between non-height traits and a given region of the genome were identified consistently when using each of the two genetic marker datasets. Eighteen MTAs between non-height traits and a given region of the genome were identified only when using the 2013 genetic marker dataset and 24 only when using the 2020 genetic marker dataset (Figure S3F). The total non-height MTAs which would likely be detectable based on allele frequency and effect size in this population of 304 sorghum lines with sufficient numbers of markers was estimated to be approximately 85 (Lincoln Index Method). GWAS with either one of the two genetic marker datasets identified only 35-43% of the estimated total number of MTAs and the combined analysis identified only 63% of the estimated total number of potentially discoverable MTAs (given sufficient marker density). Hence, increases in the number of genetic markers scored in this population would likely enable the discovery of 50-200% more MTAs when analyzing existing published trait data via single trait GWAS.

### Limited evidence for pleiotropy from conventional genome-wide association

GWAS was conducted for the complete set of 234 traits and all genetic marker data for the set of 343 accessions present in the 2020 marker dataset with the goal of testing for evidence of pleiotropy for quantitative traits. Analysis was conducted using three distinct statistical approaches to GWAS: GLM, MLM, and FarmCPU. The significant peaks identified by each of these methods are provided on FigShare (see data availability statement) for researchers interested in obtaining lists of loci controlling variation in specific traits. However, here we specifically present the results of MLM-based GWAS. With this larger population of individuals – 343 vs the 304 shared between the 2013 and 2020 datasets – a total of 56 significant peaks were identified across 43 traits. After identifying and merging associations between distinct traits within the same genomic intervals, the set of 56 significant peaks collapsed to 31 regions of the genome (Figure 3).

**Figure 3.**
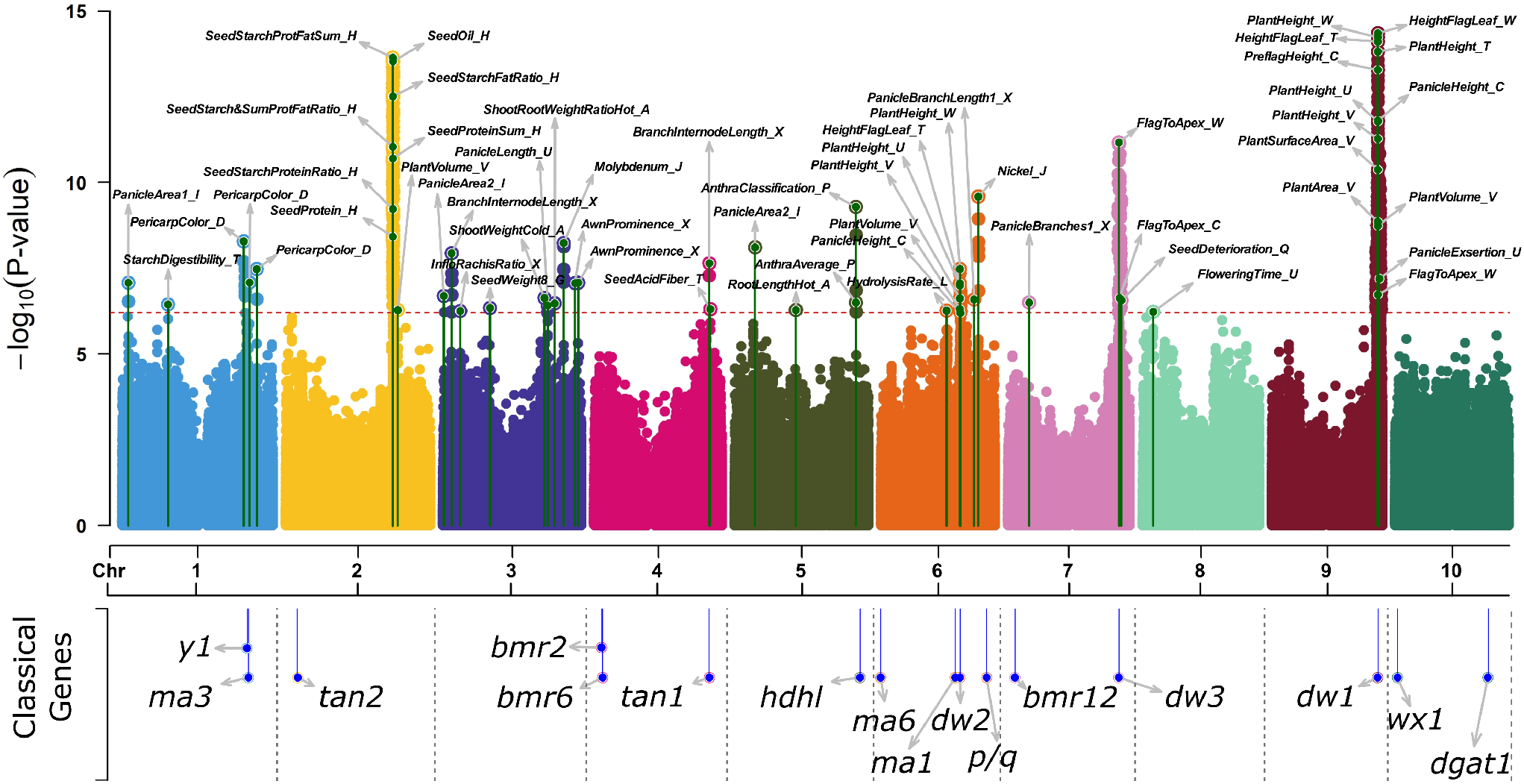
Combined Manhattan plot for GWAS using all 343 individuals genotyped in the 2020 SNP set. Combined Manhattan plot for 43 traits with at least one significant GWAS hit when analyzed using the 2020 genetic marker dataset and all 343 sorghum lines genotyped in the 2020 genetic marker dataset. Green lines topped with circles indicate the physical position and −log10 p-value for the single most significant SNP within a GWAS peak identified for a given trait. Text labels employ trait names provided in Table S1. Dashed red line indicates the cutoff for statistical significance calculated from the effective SNP number in the 2020 genetic marker dataset. Lower panel indicate positions of a set of cloned sorghum mutants, taken from^22^.

Of these 31 unique genomic regions, 25 were identified in the analysis of only a single trait dataset. Among the remaining six cases where two or more trait datasets identified signals in the same genomic regions, three were identified in the analysis of only two trait datasets. The final three intervals (on chromosomes 2, 6, and 9) were associated with 7, 6, and 13 traits, respectively. The peak on chromosome 2 was identified in GWAS for seed composition traits including oil, protein and the sums and ratios of seed oil, protein and starch and likely corresponds to the putative alpha-amylase 3 gene, previously identified in^55^. The peaks on chromosomes 6 and 9 correspond to *ma1*/*dw2*^68, 70^ and dw1^71,72^, respectively. Traits associated with these two genomic intervals include measures of both plant height and plant volume/area (Table 2).

**Table 2.**
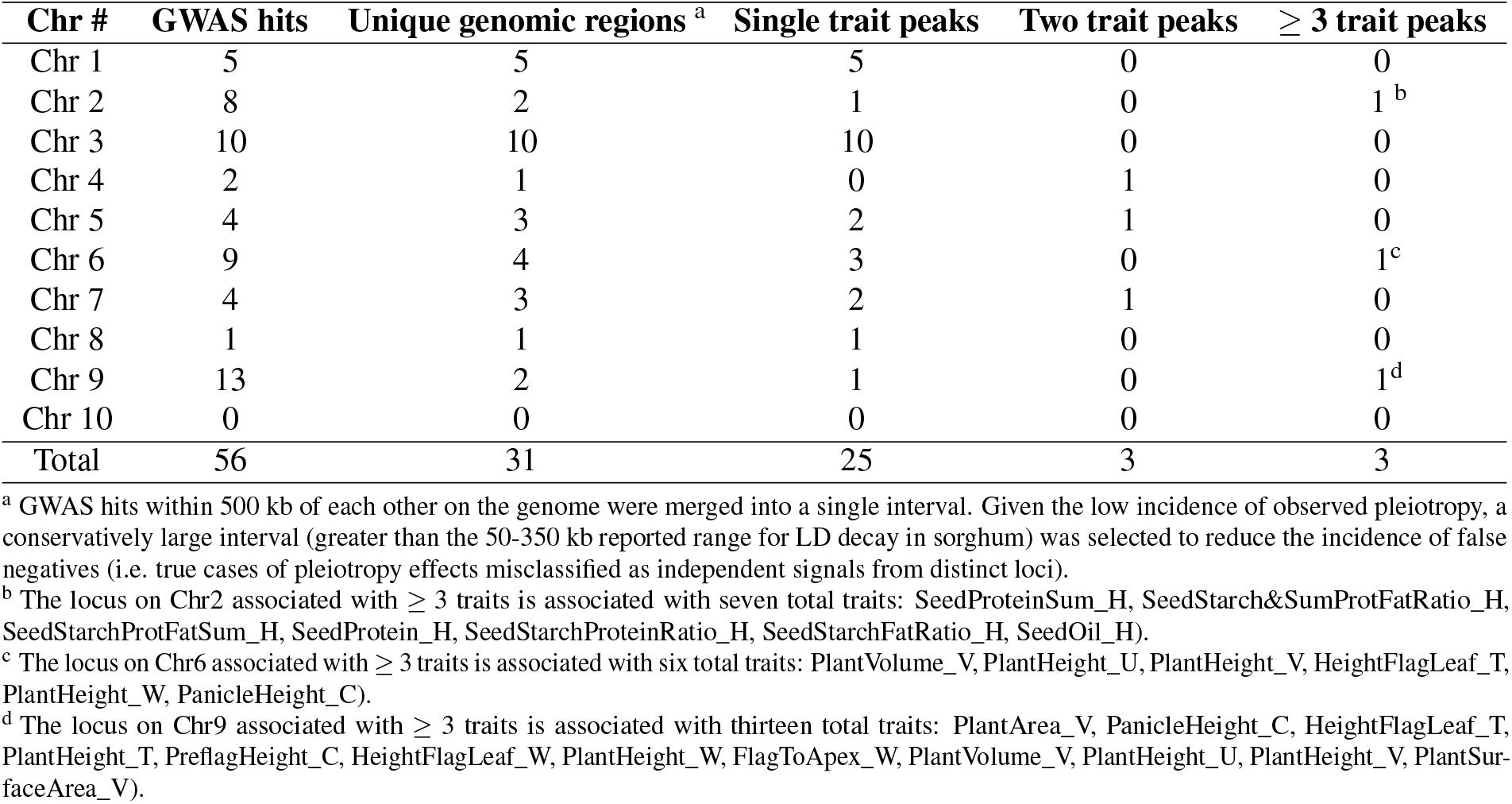
Summary of the GWAS results when data from all 343 accessions in the 2020 marker set are employed

| Chr # | GWAS hits | Unique genomic regions <sup>a</sup> | Single trait peaks | Two trait peaks | ≥ 3 trait peaks |
| --- | --- | --- | --- | --- | --- |
| Chr 1 | 5 | 5 | 5 | 0 | 0 |
| Chr 2 | 8 | 2 | 1 | 0 | 1 <sup>b</sup> |
| Chr 3 | 10 | 10 | 10 | 0 | 0 |
| Chr 4 | 2 | 1 | 0 | 1 | 0 |
| Chr 5 | 4 | 3 | 2 | 1 | 0 |
| Chr 6 | 9 | 4 | 3 | 0 | 1 <sup>c</sup> |
| Chr 7 | 4 | 3 | 2 | 1 | 0 |
| Chr 8 | 1 | 1 | 1 | 0 | 0 |
| Chr 9 | 13 | 2 | 1 | 0 | 1 <sup>d</sup> |
| Chr 10 | 0 | 0 | 0 | 0 | 0 |
| Total | 56 | 31 | 25 | 3 | 3 |

<sup>a</sup> GWAS hits within 500 kb of each other on the genome were merged into a single interval. Given the low incidence of observed pleiotropy, a conservatively large interval (greater than the 50-350 kb reported range for LD decay in sorghum) was selected to reduce the incidence of false negatives (i.e. true cases of pleiotropy effects misclassified as independent signals from distinct loci).
<sup>b</sup> The locus on Chr2 associated with ≥ 3 traits is associated with seven total traits: SeedProteinSum\_H, SeedStarch&SumProtFatRatio\_H, SeedStarchProtFatSum\_H, SeedProtein\_H, SeedStarchProteinRatio\_H, SeedStarchFatRatio\_H, SeedOil\_H).
<sup>c</sup> The locus on Chr6 associated with ≥ 3 traits is associated with six total traits: PlantVolume\_V, PlantHeight\_U, PlantHeight\_V, HeightFlagLeaf\_T, PlantHeight\_W, PanicleHeight\_C).
<sup>d</sup> The locus on Chr9 associated with ≥ 3 traits is associated with thirteen total traits: PlantArea\_V, PanicleHeight\_C, HeightFlagLeaf\_T, PlantHeight\_T, PreflagHeight\_C, HeightFlagLeaf\_W, PlantHeight\_W, FlagToApex\_W, PlantVolume\_V, PlantHeight\_U, PlantHeight\_V, PlantSurfaceArea\_V).

One interval on chromosome 7 where a single genomic interval contained MTAs for two and only two trait datasets was the result of measurement of the same trait – distance from the flag leaf to the plant apex – in two different studies conducted in different years in different locations by different research groups. An interval on chromosome 5 was associated with two traits from the same publication, which scored anthracnose resistance in two different ways^56^. A third interval on chromosome 4 was associated with both branch length in the inflorescence and acid detergent fiber within the grain. This interval on chromosome 4 may represent genuine pleiotropy or two distinct functional variations in different genes separated by ≤ 500 kilobases. The relative dearth of evidence for pleiotropy in sorghum is consistent with previous quantitative genetic investigations of pleiotropy in maize for both inflorescence architecture and leaf morphology^15,16^.

However, given the large number of false negatives expected in any individual GWAS^73^, quantifying how many intersections exist between independently conducted sets of GWAS in modestly powered populations is likely to underestimate the true extent of pleiotropy^74^. Formal multivariate GWAS approaches face difficulty scaling to large datasets (>3-5 traits)^14,75^. Hence, here we employed a multivariate adaptive shrinkage approach to the initial output from MLM based GWAS to estimate the effect of individual markers on separate trait datasets^32^. This approach provides both a test of which markers are significantly associated with phenotypic variation across the population while also estimating which specific traits a given marker had non-zero effects on, and the directions of those effects.

### Improved interpretability of trait associated loci under joint analysis

Joint analysis was conducted using MashR for 176 traits, excluding 30 traits scored on no more than 100 individuals, six binary traits and 22 ionomics trait data^46^. Standard error and effect size from MLM based GWAS were employed for this analysis. While FarmCPU has been shown to exhibit greater power to detect more total causal loci, the inclusion of identified loci as covariates means that only a single marker is identified per locus. If different markers in LD with the same causal locus were identified in FarmCPU based analysis of different traits, the pleiotropic effects of this locus would be undetectable by MashR. A set of 593 markers were identified which both exhibited an association with at least one phenotype with a local false sign rate (lfsr) <0.001, and for which the ratio of the likelihood of one or more significant phenotypic effects at a SNP to the likelihood that the SNP had only null effects was estimated to be <10^−4^, which is referred to as the Bayes factor^32^. An analog of the false discovery rate, *lfsr* requires true discoveries to be not only nonzero, but also correctly signed^76^. These 593 markers cluster together in 44 unique peaks across the sorghum genome (Figure 4). Within each multi-marker peak, the single marker with the largest Bayes factor was employed to represent the peak. The number of traits upon which markers had significantly nonzero effects ranged from only one to 141 for a peak on chromosome 6 corresponding to two known large effect loci, *dw2* and *ma1* (Figure 4A-B). The relationship between the Bayes factor assigned to a peak and the number of trait datasets with which it was associated was not straightforward (Figure 4B; S5). The single peak with the largest Bayes factor was associated with only twelve trait datasets, a number of which were independent measures of the same traits in different environments. Other peaks with comparatively modest Bayes factors were associated with modest effects on 90 traits in our datasets (Figure 4A).

**Figure 4.**
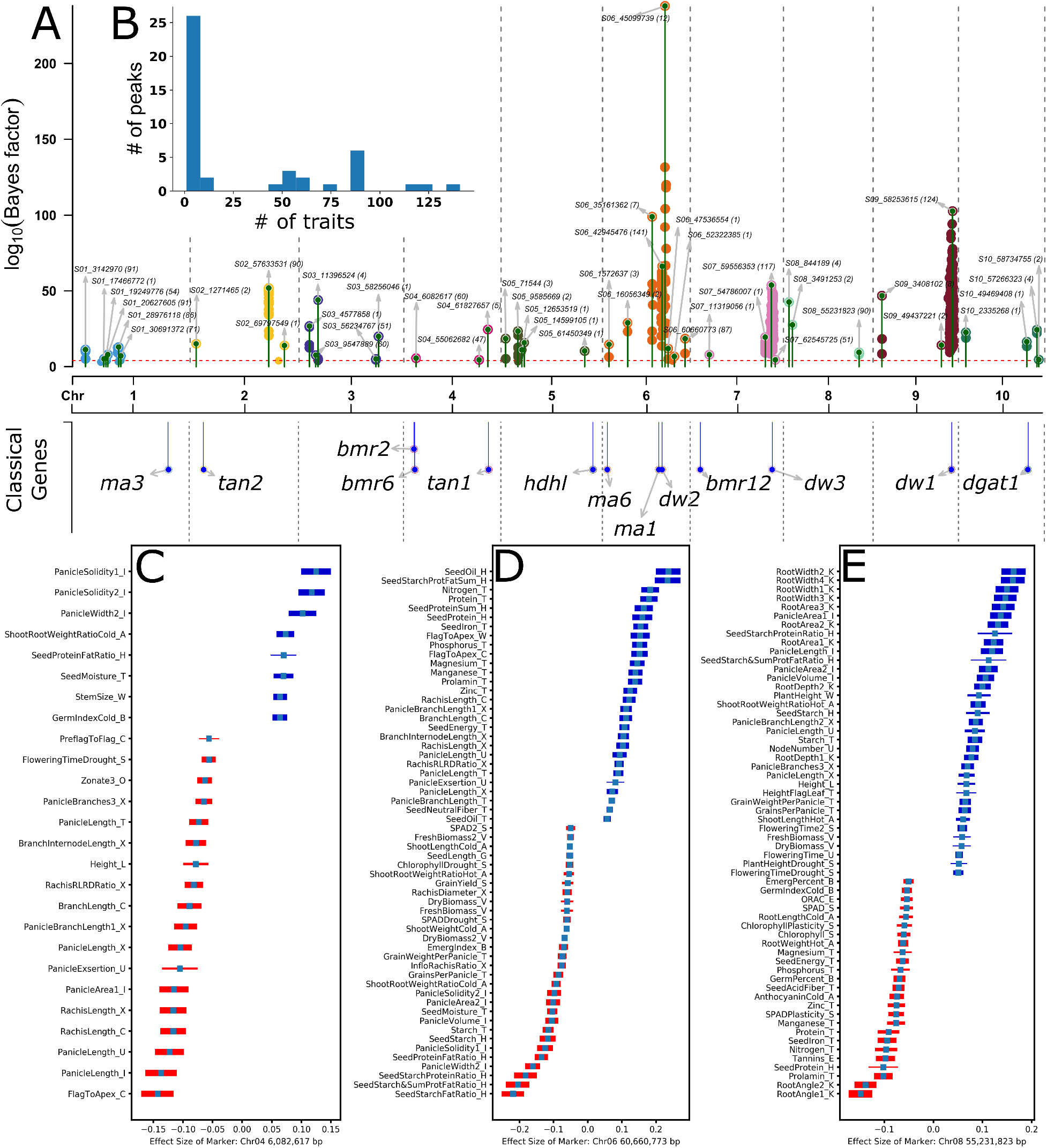
Pleiotropic analysis of SAP phenotypes. A) Markers assigned significant Bayes factor values in MashR analysis. Green lines topped with circles indicate the physical position and log10 Bayes factor for the most significant SNP within a peak identified for a pleiotropic loci. Text labels indicate the position and name of the most significant marker within each peak. The number of trait datasets significantly associated with a marker is indicated in brackets. It should be noted that trait datasets include both measurements of different traits and the same trait scored across different environments in different studies. Dashed red line indicates the cutoff for statistical significance at log10 Bayes factor of 4. B) Distribution of the number of trait datasets which were significantly associated with each unique peak. C) Distribution of effect sizes and directions of a subset of the 60 trait datasets for which the genetic marker S04_6082617 has a significant effect (*lfsr* <0.001). To aid readability, only the subset of trait datasets where the effect size is >0.05 or <0.05 are shown. Bar thickness is proportional to the relative statistical significance of each effect. D) Distribution of effect sizes and directions for a subset of the 87 trait datasets for which the genetic marker S06_60660773 has significant effects. Cutoffs for visualization are the same as applied for panel C. E) Distribution of effect sizes and directions for a subset of the 90 trait datasets for which the genetic marker (S08_55231823) has significant effects. Cutoffs for visualization are the same as applied for panel C and D.

Multi-trait analysis was able to recover a number of known pleiotropic features for large effect loci segregating in the population. In addition to plant height, the peak at *dw2* was associated with multiple matrices of root size/area, panicle length, plant surface area and seed weight (Figure S4A). The effects of *dw2* on panicle length, seed weight and leaf area have been previously reported,^77,78^ while the reductions in multiple metrics of root size/area associated with the dwarfing allele of *dw2* had not. The apparent impact of *dw2* on root phenotypes, suggests that the gene may play equivalent roles in determining size of below and above ground plant organs. Another plant height related gene, *dw3* was previously known to have effects on grain yield^79^, leaf angle^80^, biomass^81^, internode length^25^, stem diameter^82^, panicle exsertion^47^ and panicle architecture^83^. Here, statistically significant links were also observed between *dw3* and variation in plant biomass, leaf angle, stem size/stem diameter, grain yield plasticity, internode length, panicle exsertion and panicle architecture (panicle length, width and area) (Figure S4B). In addition to plant height, *dw1* is also known to alter biomass and biomass associated traits^57^, internode length and lodging resistance in sorghum^71,72^. The set of traits assembled here did not include measurements of either internode length or lodging resistance across the sorghum association panel so it was not possible to assess whether these known pleiotropic effects of *dw1* on these traits were recovered. However, the peak associated with *dw1* in chromosome 9 was significantly linked to variation in above ground biomass traits (directly measured biomass, plant surface area), as well as multiple metrics for root size/area (Figure S4C). In contrast to *dw3* and similar to dw2, *dw1* may play equivalent roles in determining organ size for both below ground and above ground plant organ systems.

Multi-trait analysis also recovered a number of novel signals across the genome. The pairing and direction of effect sizes for these traits enables greater interpretability of the resulting MTAs. In some cases, these are straightforward trade-offs. The trait associated marker located at 6.08 MB on chromosome 4 one allele is associated with longer panicles, but these panicles are also narrower and less dense. The other allele present at this locus produces shorter, fatter, and denser panicles, resulting in increases in seed moisture levels at harvest (Figure 4C). A number of other multiple trait associations identified were also consistent known trade-offs in plant growth and development. A trait associated marker located at 60.66 MB on chromosome 6 is associated with increases in seed oil and protein content, and many important micronutrients. However, the same allele is also associated with decrease in panicle volume and solidity (high throughput phenotyping) and decreases in grains per panicle and grain weight per panicle (conventional phenotyping) (Figure 4D). Multiple trait associations can also reveal explanations for potential associations which would otherwise be potential breeding targets. For example, improving root architecture has been proposed as a target for enhancing drought tolerance or nutrient uptake^84^. A marker on chromosome 8 located at 55.23 MB had large effects on multiple root traits including greater root width, larger total root area, and a smaller root angle (Figure 4E). In isolation this might appear a promising target for root-based breeding. However, multiple trait analysis also identified that this allele is associated with delays in flowering time and increases in total node numbers. These results suggest that the observed increases in root extent and root area may be an indirect result of delayed vegetative to reproductive transition.

## Discussion

The SAP has been widely adopted and proven to be a long-lasting resource for the sorghum genetics community. The syntenic conservation of GWAS hits between sorghum and maize means the SAP also provides information on gene function in maize^63^. In the interval between the start of the analyses in this paper and submission for publication, at least five additional studies employing this population have been posted online including studies of provitamin A^85^, geospatial association with parasitic plants^86^, resistance to different fungal sources of grain mold^87^, genetic determinants of the root associated microbiome^88^ and herbicide resistance^89^. Our results suggest that simply increasing the marker density of the SAP – and similar community association populations – may more than double the number of true positive MTAs detection in future studies with the same population. Care should be taken to record and disseminate accession specific trait measurements from GWAS in ways that facilitate future reanalyses as additional genetic marker datasets become available. In our study, we identified 40 papers which included the collection of one or more new trait datasets from the SAP. Through data curation and annotation, we have increased the proportion of papers from 50% to 64% where traits have been publicly released with IDs which can be associated back to genetic marker data, facilitating reuse and reanalysis (Table 1). However, adopting both community norms that emphasize the need to store and disseminate trait datasets, as well as developing a central repository for sorghum phenotypic data would likely increase the proportion of trait datasets generated by the sorghum genetics community, which will continue to contribute to new discoveries and understanding in years to come. Similarly, it will be important to maintain and distribute seed of the SAP to lower barriers to entry into sorghum quantitative genetics for new research groups and to avoid the risk of failed or misleading results due to lines that are swapped or duplicated. Seed is currently maintained and distributed by the USDA NPGS, however resource constraints at NPGS can limit how often the lines of this panel can be increased. Informal lab to lab distribution has acted as a fallback source of seed. However, in the long term this approach runs the risk of propagating swaps, labelling errors or pollen contamination, reducing the comparability of data generated by different research groups with different germplasm sources. An analysis of published RNA-seq data labeled as coming from the maize reference inbred B73 found at least three distinct clades of genetically distinct B73 accessions and that relationships between these samples recapitulated advisor-advisee relationships^90^. In arabidopsis, a widely used commercial source for the reference inbred Col0 was found to contain substantial introgressions of non Col0 origin^91^. Storage and dissemination of trait data and associated metadata for future studies will aid both in detecting new associations as genetic marker datasets increase in density, and in expanding our knowledge of pleiotropy, as shown here. Additionally, this community-based strategy will facilitate the development and validation of predictions from empirical crop growth models, and the investigation of the genetic basis of phenotypic plasticity and genotype by environment interactions.

Strong selection will often act on rare, large effect, and pleiotropic loci^92^. Here, loci identified in a joint analysis of 176 trait datasets tended to fall into one of two categories, either showing associations with large (>40) number of trait datasets or small (<10) trait datasets, with these datasets often representing measures of the same trait in multiple environments or multiple distinct but highly correlated traits (Figure 4B). This pattern does not appear to be an artifact introduced by variation in statistical power as a result of either effect size or allele frequency as both loci associated with many traits or associated with only several traits include examples with both high and low Bayes factor values (Figure S5). This stands in contrast to studies in the related species of maize where little evidence has been found for pleiotropic quantitative genetic loci segregating in populations^15,18^, but this difference should be interpreted cautiously until and unless similar wide-scale multivariate analyses are conducted in maize association populations, given the differences in both methodological approaches and patterns of linkage disequilibrium. An analysis of historical yield data in common bean employing MashR also identified two genomic intervals associated with pleiotropic effects on different phenotypes^93^. If the difference in the prevalence of pleiotropy between maize and sorghum continues to be observed in additional studies, it may reflect the distinct histories of both maize and sorghum in temperate North America, and the distinct histories of widely used association panels in each species. The first reports of sorghum cultivation in the southeastern United States date to either 1838 or 1855, likely as the result of introduction from the Caribbean^94^. Two temperate adapted strains of sorghum were introduced into the Great Plains approximately 150 years ago followed by rapid selection by farmers for earlier flowering and shorter stature^95^. Temperate maize in the United States is much older with adaptation to temperate highlands occurring over an approximate 2,000 year period starting 4,000 years ago, allowing for more gradual selection and therefore less likely to capture pleiotropic loci^96^. Similarly, many of the lines in the SAP trace their origin to a conversion process where genes needed for temperate adaptation were introgressed through multiple generations of strong phenotypic selection^20,23^, while both the most widely employed maize association panel and the maize nested association panel were assembled from lines already adapted to the temperate United States^6,97^.

As high throughput phenotyping approaches become more widely adopted, direct measurements of pleiotropy may become more feasible. Once sensor datasets are collected (e.g. RGB images, LIDAR point clouds, hyperspectral data cubes) and algorithms for numerically quantifying specific traits are developed, the additional cost extracting measurements of non-target traits from the same sensor data is minimal. A better understanding of pleiotropic relationships for specific loci and across groups of plant traits may aid significantly in reducing inadvertent selection and prioritizing candidate loci for introgression into elite germplasm for sorghum and related species. A greater understanding of potential pleiotropic effects may help to prioritize which off target phenotypic effects should be tested for either when evaluating natural variants or when generating gene edits.

## Methods

### Genetic Marker Datasets

A set of 265,487 SNPs generated using conventional genotyping by sequencing and aligned to version 1 of the BTx623 sorghum reference genome were downloaded from http://people.beocat.ksu.edu/gpmorris/sorghum_GBS_data/readme.txt/^30^ (the “2013 dataset”). A set of 569,305 SNPs generated using a modified genotyping by sequencing approach and aligned to version 3 of the sorghum genome was obtained from FigShare (https://doi.org/10.6084/m9.figshare.11462469.v5)^31^ (the “2020 dataset”). In each dataset missing data points were imputed using Beagle (v4.1) with the sliding windows set individually for each chromosome to capture 10% of all markers on that chromosome and overlap windows set to capture 2% of call markers on that chromosome^98^. After imputation, marker sets were filtered by removing markers with minor allele frequencies of less than 5% among the set of genotypes employed for a given analysis. This filtering criteria resulted in a set of 107,751 markers scored across 304 lines for the 2013 dataset and a set of 257,882 markers scored across the same 304 lines for the 2020 dataset. Filtering using the same parameters across all 343 lines included in the 2020 dataset produced a set of 256,695 markers.

### Trait Datasets

A total of 234 trait datasets scored across all or subsets of the sorghum SAP were employed in this study. One hundred and ninety of these trait datasets were drawn from published sources as described in Table S1. An additional 12 phenotypes were generated using sums or ratios of published trait datasets. The traits and formulas used to generate these 12 phenotypes are provided in Table S2. The remaining 32 trait datasets employed in this study were previously unpublished datasets collected at either the University of Nebraska-Lincoln in Nebraska (19 datasets) or Clemson University in South Carolina (13 datasets).

#### Nebraska Trait Collection

A single replicate of the sorghum association panel was grown near Mead, NE in 2016 and 2017. Plant height to inflorescence, plant height to flag leaf, leaf angle (3rd leaf), stem diameter (between the 3rd and 4th leaf) and node number were measured from a representative plant at reproductive maturity in 2016. A ratio of plant height to inflorescence/plant height to flag leaf was also calculated (Table S2). Inflorescence architecture traits were measured using two representative plants at maturity in 2016, 2017 and included inflorescence length, rachis length, rachis diameter, number of primary branches, length of primary branches at the bottom third of the inflorescence, length of primary branches at the top third of the inflorescence, number of secondary branches on a primary branch, number of third-order branches on a secondary branch, first internode length on a primary branch, prominent awns (binary trait), and prominent glumes (binary trait). During 2017 one additional trait, infertility (scored on a scale from 1-4), was also collected (Figure S6). Two ratios were also calculated from each year in this dataset: inflorescence length/rachis length, rachis length/rachis diameter (Table S2). Best linear unbiased predictors (BLUPs) for each phenotype were calculated by fitting a linear mixed model using R package lme4^99^ with genotype, year and genotype by year variables fit as random for traits with multiple years of data and only genotypes fit as random variable for the traits with data from only one year.

#### South Carolina Trait Collection

The sorghum association panel was grown near Florence, South Carolina in 2013, 2014, and 2017. In each year two replicates per line were grown in a 2x replicated completely randomized design utilizing two row yield plots. Trait datasets collected at Clemson University included two flowering time related traits, measured in all three years: days to anthesis and grain fill duration (days to maturity - days to anthesis). Two plant height traits were measured: plant height from ground/plant base to panicle apex and flag leaf height. Six reproductive traits were measured: number of grains per primary panicle and grain yield per primary panicle, measured in all three years, glume tenacity (0-5 visual rating), primary panicle branch length, panicle length, and exsertion in 2017. Five biochemical traits: magnesium (% dry basis), manganese (ppm), nitrogen (mg), phosphorus (% dry basis), and zinc (ppm) measured from ground grain samples in 2013 and 2014 using near-infrared spectroscopy^48^. Thirteen seed traits were determined from these experiments: percent moisture, 1000-grain weight, percent of dry mass which was acid detergent fiber and percent of dry mass which was neutral detergent fiber, percent of starch which is amylose, percent of dry grain weight which was oil, protein, or starch, in vitro starch digestibility, gross energy per gram (calories/gram), iron (ppm), prolamin as a percentage of dry weight, and seed density (grams per milliliter). All except 1000-grain weight and seed density were measured using NIR and were evaluated in all three years. All phenotypes were measured in all three years except seed density which was measured only in 2017. For each trait a linear mixed model was fit using the R package lme4^99^ with genotype, year, genotype by year and replication nested in a year were fit as random for traits from 2013, 2014 and 2017 combined and the genotype and replication fit as random for the traits collected only during 2017, and the resulting phenotypic BLUPs were employed for further analysis. Of the 28 traits included in this study, 15 were published in part or in totality^45,48,100^ and 13 are previously unpublished data.

#### Trait Data Normalization and Heritability Calculation

With the exception of six binary traits (AnthraClassification_P, PericarpColor_D, Tannins_D, Tannins_F, AwnProminence_U and GlumProminence_U) each trait dataset was normalized before analysis using the R package, bestNormalize version 1.4.3^101^. The function bestNormalize in the bestNormalize package performs various/suite of normalization transformations, such as Lambert W x F, BoxCox, YeoJohnson, Ordered Quantile, etc and then select one optimal transformation for each dataset based on minimizing the Pearson P statistic, a test for normality. Subsequent to normalization, values which were more than 1.5 times the interquartile range below the 25^th^ percentile of normalized trait values or more than 1.5 times the interquartile range above the 75^th^ percentile of normalized trait values were converted to missing data. In order to determine the proportion of genetic variation explainable by genetic factors, marker-based estimate of narrow-sense heritability was generated using the R-package sommer (v4.1.1), the reported values for each line for each trait, and the 2020 genetic marker dataset^102^.

### Genome-wide Association Analyses

GWAS was performed independently on each of 234 trait datasets. The number and identity of lines evaluated for various traits varied both across and within studies (Table S1). For each trait dataset, analyzed with each of the two genetic marker datasets, genetic markers were separately filtered to remove those markers with a minor allele frequency of <5% among individuals with recorded values for the target trait.

The resulting marker sets were employed for GWAS analysis using two single-locus models; GLM (PC), MLM (PC+K)^103^ and one multi-locus model; FarmCPU^104^. For all three models, the implementations used were those included in the R package rMVP (v1.0.1)^105^. For the GLM model, the first three principal components (PCs) were fit as covariates in order to control for the population structure. In case of MLM model, in addition to the first three PCs, a kinship matrix was also integrated in the model for association analysis. The kinship matrix representing the relationship among individuals used in the MLM model was calculated using VanRaden method^106^ within the rMVP package. The multi-locus mixed linear model; FarmCPU, was run with the first three PCs as covariates and the kinship matrix calculated internally by the FarmCPU algorithm fitted as random effects. FarmCPU was run using maxLoop = 10, the method for selecting most appropriate bins was run with the MVP option, method.bin = “FaST-LMM^107^. The method of variance components analysis, vc.methods was set to “GEMMA” in the association analysis^108^.

Bonferroni corrections were applied based on the effective number of independent markers in each genetic marker dataset. Effective SNP numbers were calculated for each dataset using the GEC (v0.2) software package with default parameter settings^109^. Statistical significance thresholds were set to 10^−5.958^ (0.05/48,488) and 10^−6.154^ (0.05/82,5223) for 2013 and 2020 genetic marker datasets respectively when SNPs were first filtered based on their minor allele frequencies in the set of 304 sorghum varieties shared between the two datasets. The statistical significance threshold for the 2020 genetic marker dataset when markers were filtered based on their frequency in the complete set of 343 lines genotyped in that dataset was 10^−6.1379^ (0.05/82143). Trait values were available for different subsets of lines for different trait datasets. While markers were further filtered to remove low frequency markers in subset populations as described above, the same three p-value cutoffs for statistical significance were employed for all GWAS results to provide consistency across analyses. Manhattan plots were created using the CMplot R package (v3.6.2)^110^. When multiple genetic markers showed statistically significant associations after correction for multiple testing, markers on the same chromosome and separated by no more than 1 MB were merged into a single peak. A custom python script (CallPeaksBatch.py) was employed to merge nearby SNPs into peaks and identify “summit” SNPs for each peak^111^.

### Multivariate Gene-Trait Analyses

For multivariate analysis 176 traits were chosen by excluding 30 traits collected in Brazil, six binary traits and 22 ionomics trait data^46^. Estimated effect size and standard error for each of the markers in each of these 176 traits was extracted from the initial output of rMVP using the results of analysis conducted using the MLM model. First, a subset of strong signals were chosen by running a condition-by-condition analysis using ash with package ashr/v2.2-47 using the most recent code revision available on github as of 9/8/2020^112^. A second control set of estimated effect size and standard error values for a set of 90,000 markers which were not statistically significantly associated with any of the 176 traits were also extracted to aid the mash model in learning the patterns of covariance between SNPs and each phenotype in order to produce improved effect estimates of the SNPs chosen from condition-by-condition analysis. These datasets were analyzed using mashr/v0.2.40 using the most recent code revision available on github as of 9/8/2020^32^. Following the recommendations of the MashR documentation, canonical and data-driven covariance matrices were computed. The canonical covariance matrix was calculated using the MashR function “cov_canonical”. The data-driven covariance matrix was calculated by using the function “cov_pca”. A mash model was fit using both the covariance matrices. The posterior summaries were computed for each SNP in the strong sub set, choosen from condition-by-condition analysis for each phenotype. Further, the Bayes factor extracted from mash output with CBDgenomics R package93 was used to determine if a given SNP has significant phenotypic effect. Threshold of local false sign rate (lfsr) <0.001 was used to determine the number of traits associated to a given SNP^76^. Peaks were called by merging individual significant SNPs which were separated by less than 500 kilobases into peaks and selecting the single SNP within each peak with the latest Bayes factor as representative of that peak. The code used to perform this merging has been deposited on github^111^. Visualizations of mashr results were generated using the R package CMplot (v3.6.2)110 (panel A).

## Acknowledgments

This research was supported by the Office of Science (BER), U.S. Department of Energy, Grant no. DE-SC0020355 to JCS and BS, the Foundation for Food and Agriculture Research (ID: 602757) and the US Department of Energy Advanced Research Projects Agency-Energy (ARPA-E) under Contract No. 18/CJ000/01/0 and the National Science Foundation under grant OIA-1826781 to JCS and a Research Council Faculty seed grant to BS. This project was completed utilizing the Holland Computing Center of the University of Nebraska, which receives support from the Nebraska Research Initiative.

The authors thank Melinda Yerka, Jake Ziggafoos, and undergraduates, Tyler Beam, Kate Birchmeier, Alex Cox, Lexis Funk, Vanessa Funk, Jacob Givens, Courtney Krsnak, Xiangdong Liu, Matthew Lennon, Matthew Myers, and Thomas Stefaniak for assistance in plant care and phenotypic scoring.

## Data Availability Statement

All genotype data used in this study was drawn from published sources. Combined phenotypic data on both previously published and unpublished phenotypes used in this study are provided on FigShare doi: 10.6084/m9.figshare.13143389. Detailed results from GWAS are also provided on FigShare.

## Author contributions statement

RVM, MG and JCS conceived of the study. AD, SS, REB, MGSF, PSS, BS and SK conducted the experiments and collected the data. RVM, MG, and CM analyzed the data. RVM and JCS wrote the manuscript. All authors read and approved the final manuscript.

---

**Figure S1.**
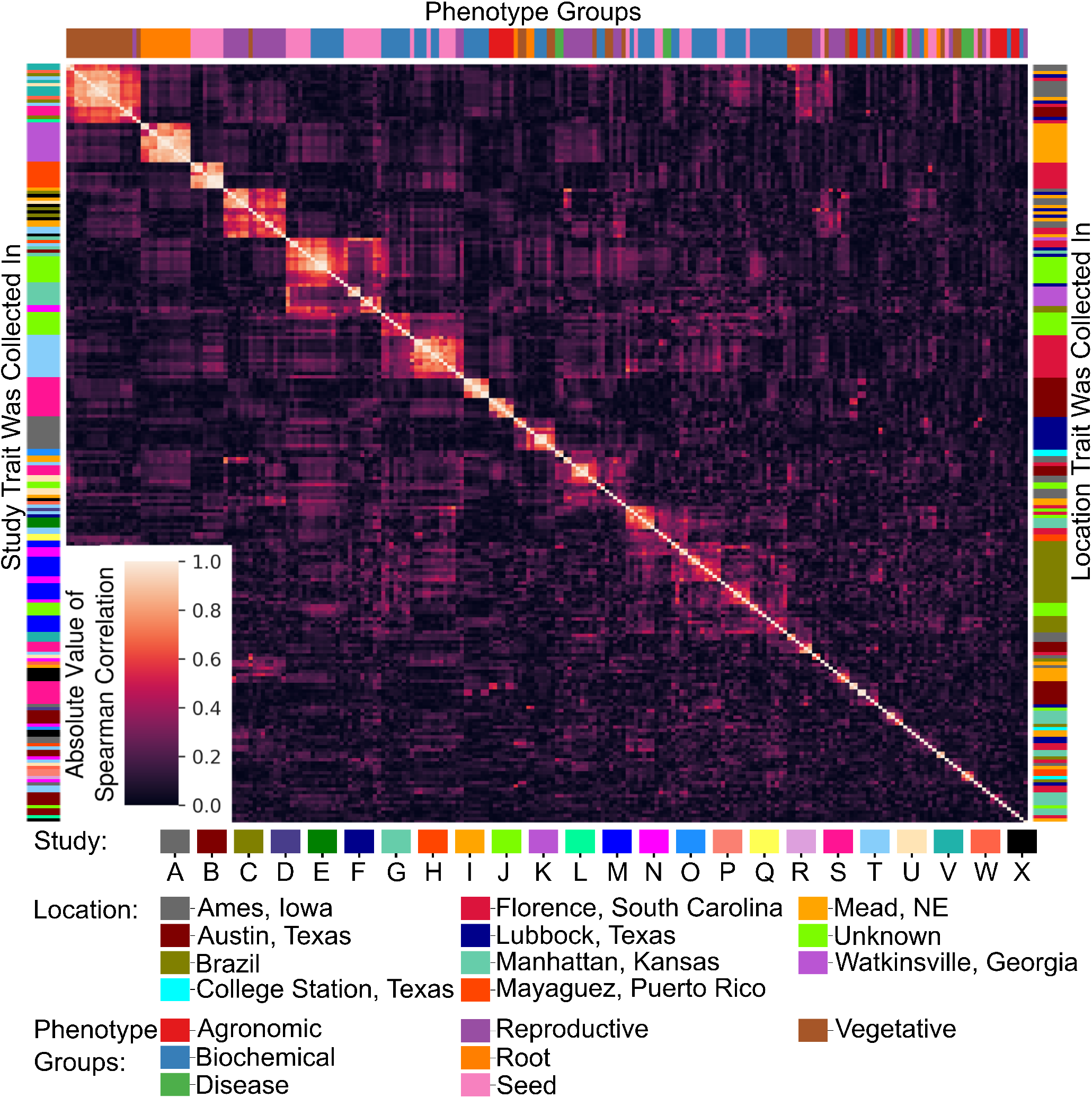
Correlations among the 234 trait datasets analyzed in this study. Trait datasets clustered based upon absolute spearman correlation value. Phenotype classes indicated with color bar on top x-axis with colors corresponding to the color key for phenotype classes provided in Figure 1 and Table S1. The identity of the study individual trait datasets were collected from, is indicated with color bar on left y-axis. Distinct colors were assigned to each of 24 studies listed in Table S1. The identity of the location where trait datasets were collected from, is indicated with color bar on right y-axis. Distinct colors were assigned to each of 24 studies listed in Table S1.

**Figure S2.**
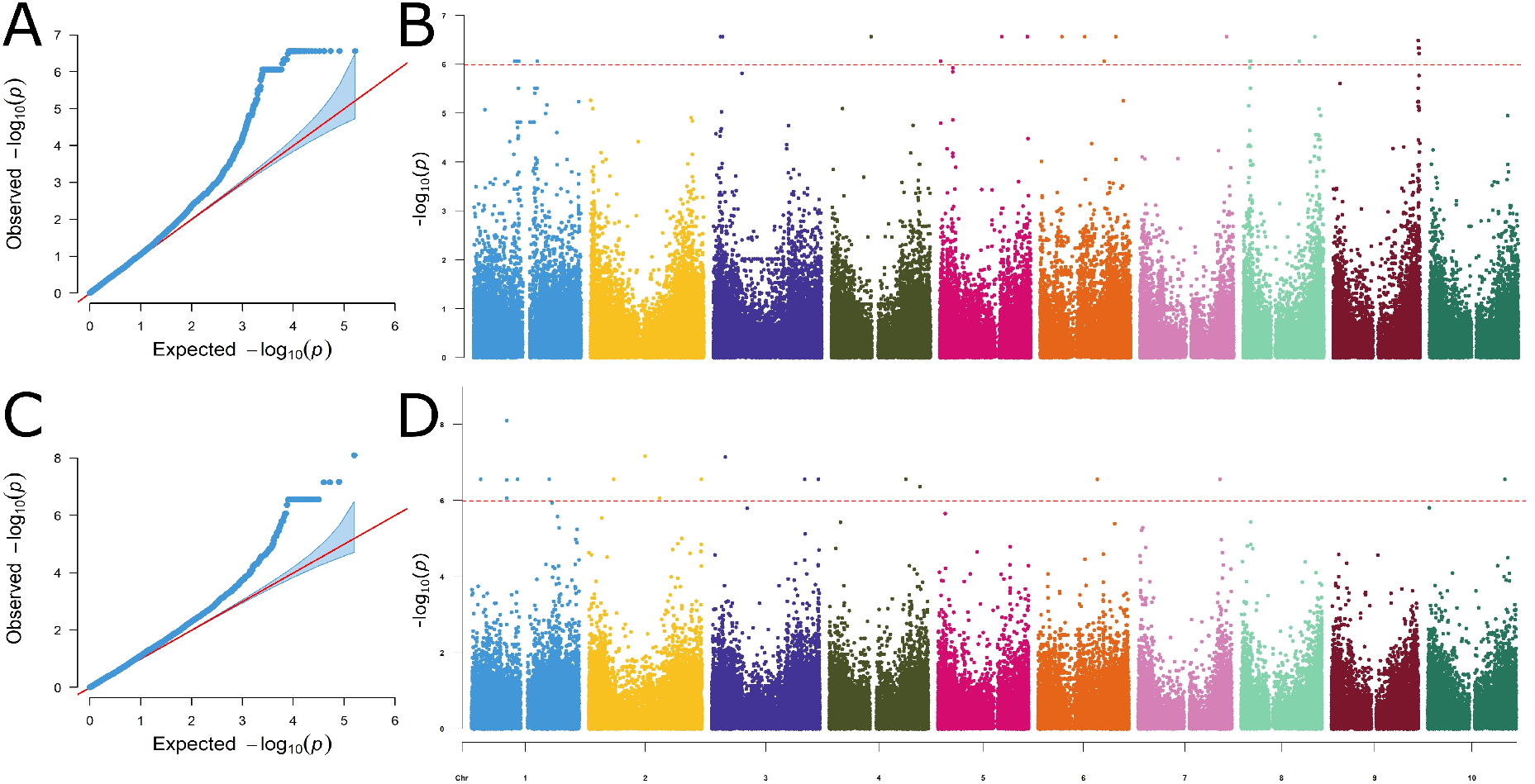
Poor MLM GWAS results for two traits which were ultimately excluded from comparative analyses. A) QQ plot for MLM based GWAS using the trait dataset SeedIron_Q and the 2013 marker set. A large number of markers have been assigned the same p-value. B) Manhattan plot for the same GWAS analysis shown in panel A. Markers with identical, and statistically significant, p-values are distributed across chromosomes 3, 4, 5, 6, 7, and 8. C) Q plot for MLM based GWAS using the trait dataset Selenium_J. A large number of markers have been assigned the same p-value. D) Manhattan plot for the same GWAS analysis shown in panel C. Markers with identical, and statistically significant, p-values are distributed across chromosomes 1, 2, 3, 4, 6, 7, and 10.

**Figure S3.**
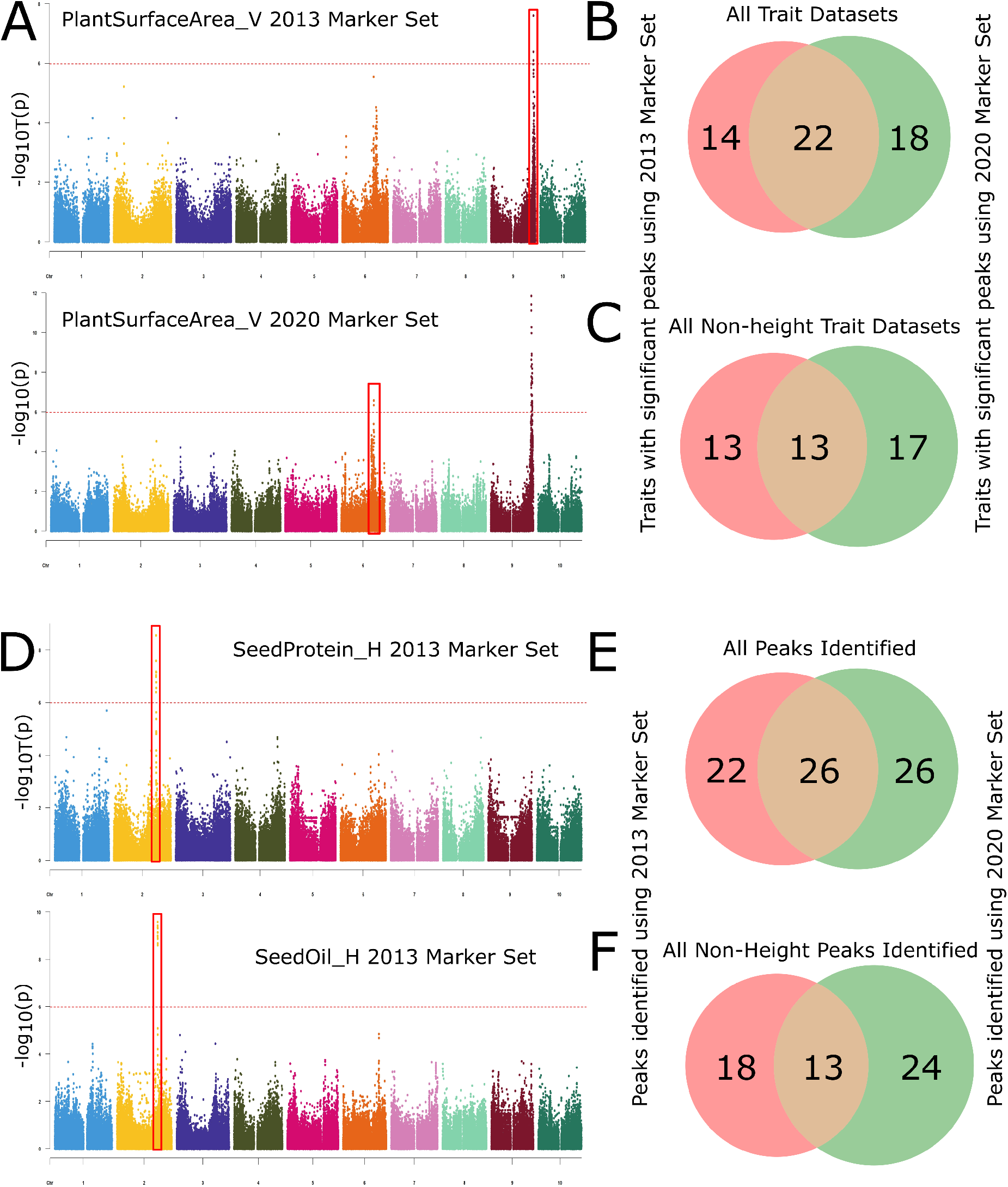
Traits exhibiting number of significant peaks as well as number of peaks identified when the same trait dataset was analyzed with each of the two genetic marker datasets. A) Example of shared traits when same phenotype is analyzed with 2013 and 2020 marker set. B) Number of traits exhibiting significant peaks considering all trait datasets when analyzed using 2013 and 2020 marker set. C) Number of traits exhibiting significant peaks for all non-height trait peaks when analyzed using 2013 and 2020 marker set. D) Example of shared peaks when two different phenotypes are analyzed with 2013 and 2020 marker set. E) Number of significant peaks identified when all trait datasets where analyzed using 2013 and 2020 marker set. F) Number of non-height significant peaks identified when all trait datasets where analyzed using 2013 and 2020 marker set.

**Figure S4.**
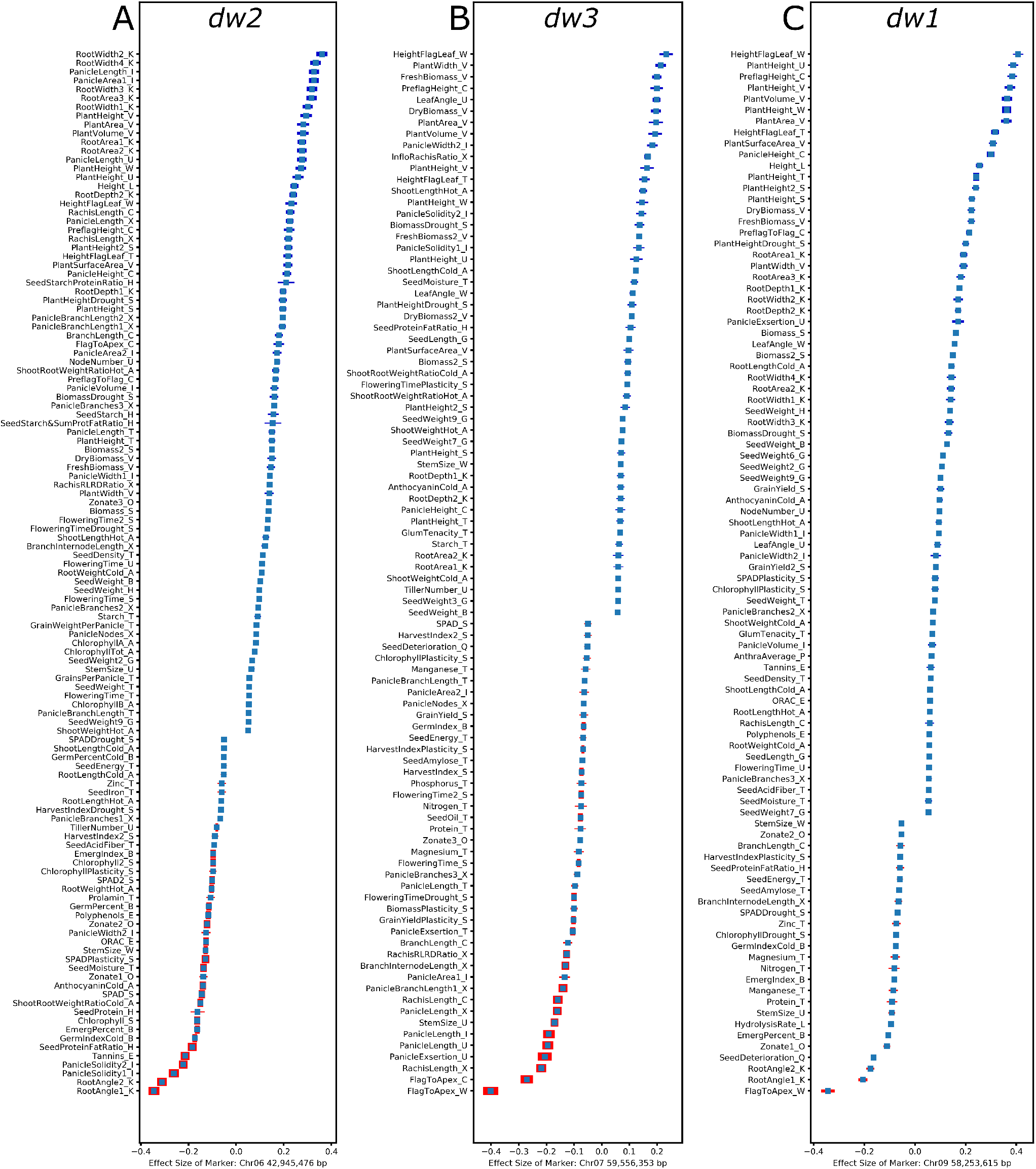
Forest plots of non-zero effect size of dwarf genes. A) Distribution of effect sizes and directions for a subset of the 141 traits for which the genetic marker S06_42945476 co-localized with *dw2* gene has a significant effect (*lfsr* <0.001). To aid readability, only the subset of traits where the effect size is >0.05 or <0.05 are shown. Thickness of bars is proportional to the relative statistical significance of each effect, with the most significant associations represented by the thickest bars and the less significant associations represented by the thinnest bars. B) Distribution of effect sizes and directions for a subset of the 117 traits for which the genetic marker S07_59556353 co-localized with *dw3* gene has significant effects. Cutoffs for visualization are the same as applied for panel A. C) Distribution of effect sizes and directions for a subset of the 124 traits for which the genetic marker S09_58253615 co-localized with *dw1* gene has significant effects. Cutoffs for visualization are the same as applied for panel A and B.

**Figure S5.**
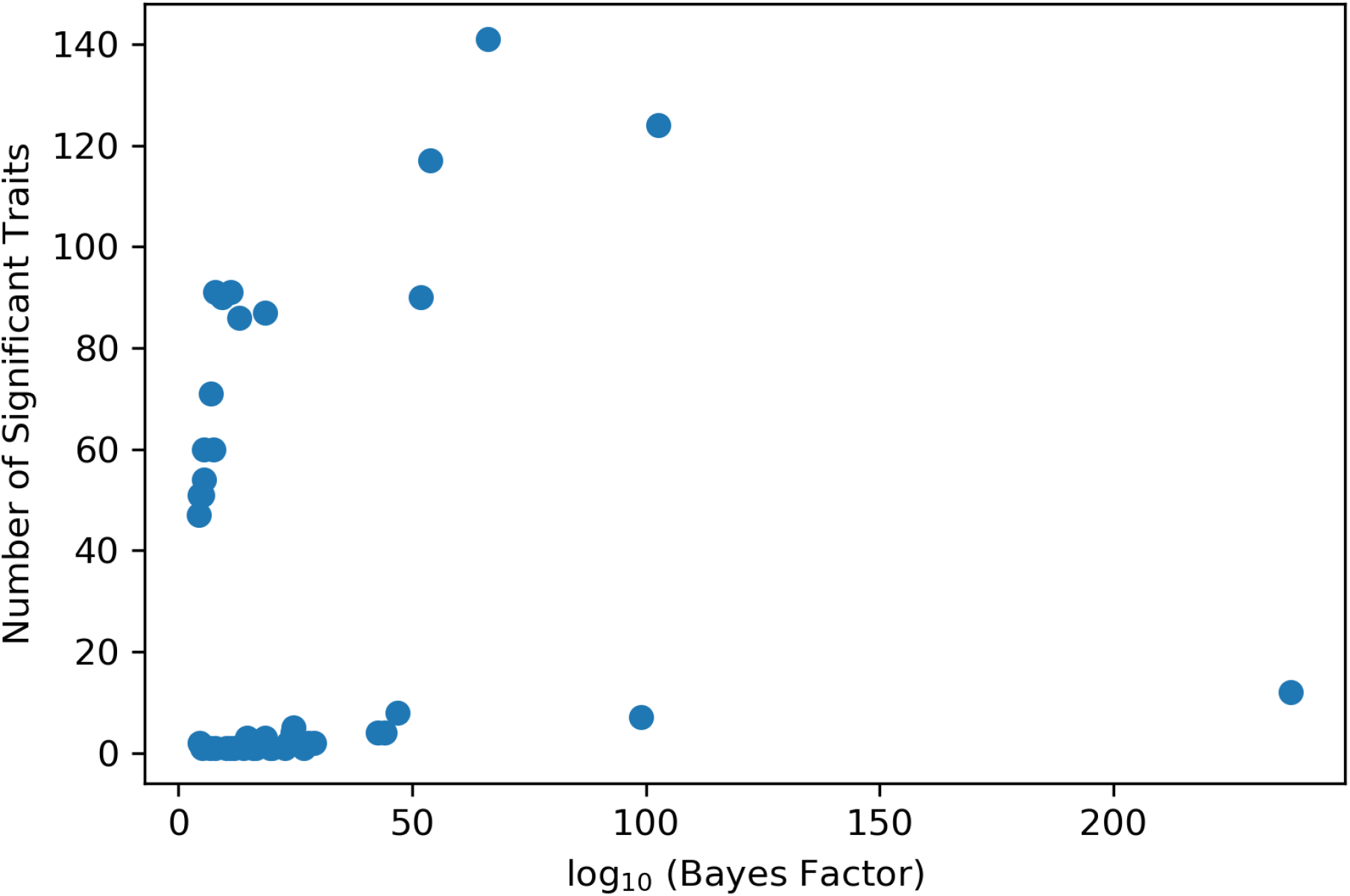
Relationship between Bayes factor and number of significantly associated traits for “summit” markers in each peak identified in Figure 4.

**Figure S6.**
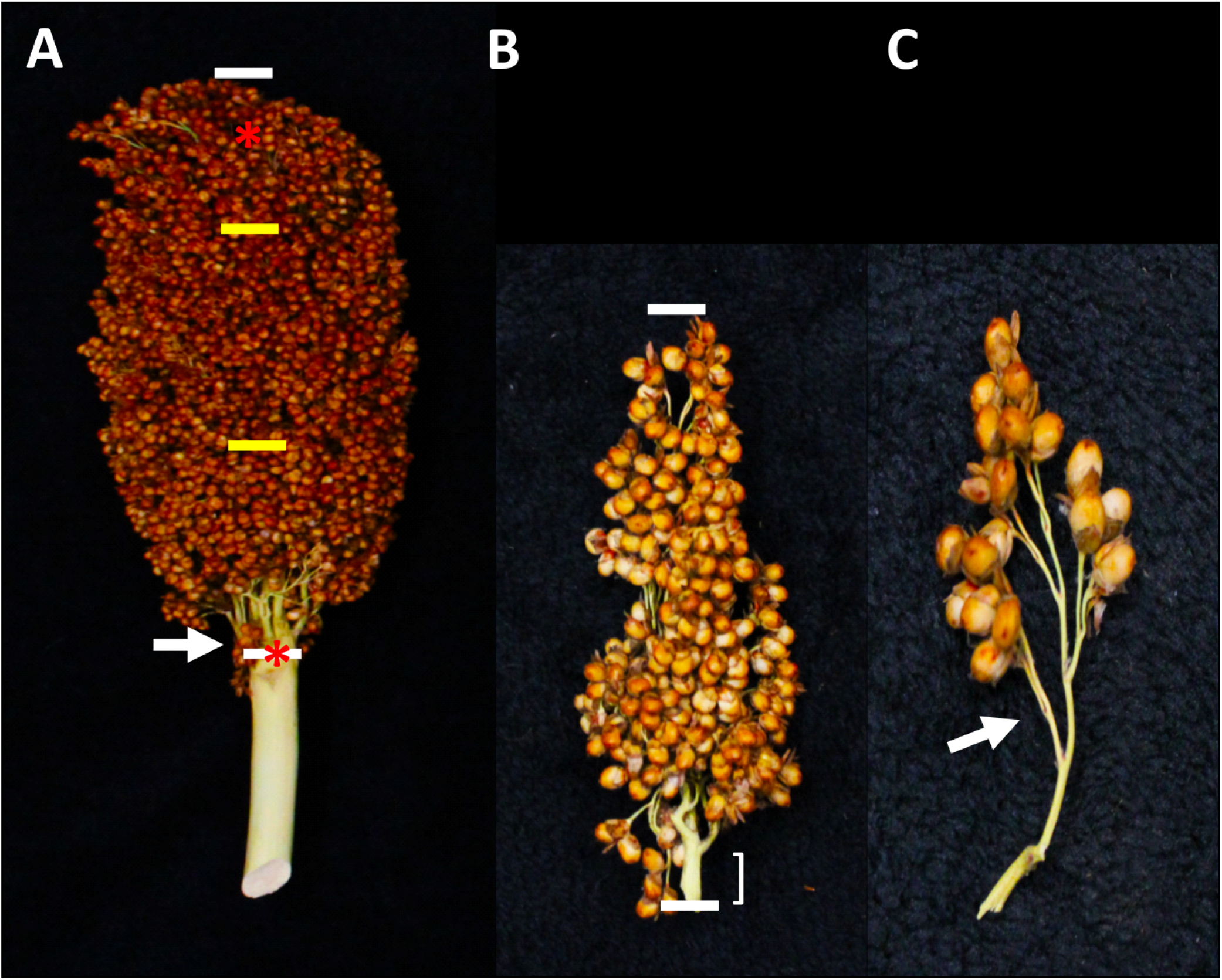
Phenotyping of Inflorescence Architecture. A) Total inflorescence length was measured from the bottom branch to the top of the inflorescence (white bars). The rachis length was also measured once all primary branches were removed and counted (asterisks). The rachis diameter was measured using digital calipers just above the removal point of the basal primary branch (arrow). A primary branch was removed and measured from the top one-third and the bottom one-third of the inflorescence, as indicated by the yellow bars. B) Primary branches were measured from the bottom of the branch to the tip of the branch (barred). The internode length was measured from the bottom of the branch to the bottom of the first secondary branch (bracketed). C) A secondary branch was also removed and the number of tertiary branches counted along the length of this branch (example indicated with an arrow).

**Table S1. Names and properties of individual trait datasets.** Provided as included excel file.

**Table S2.**
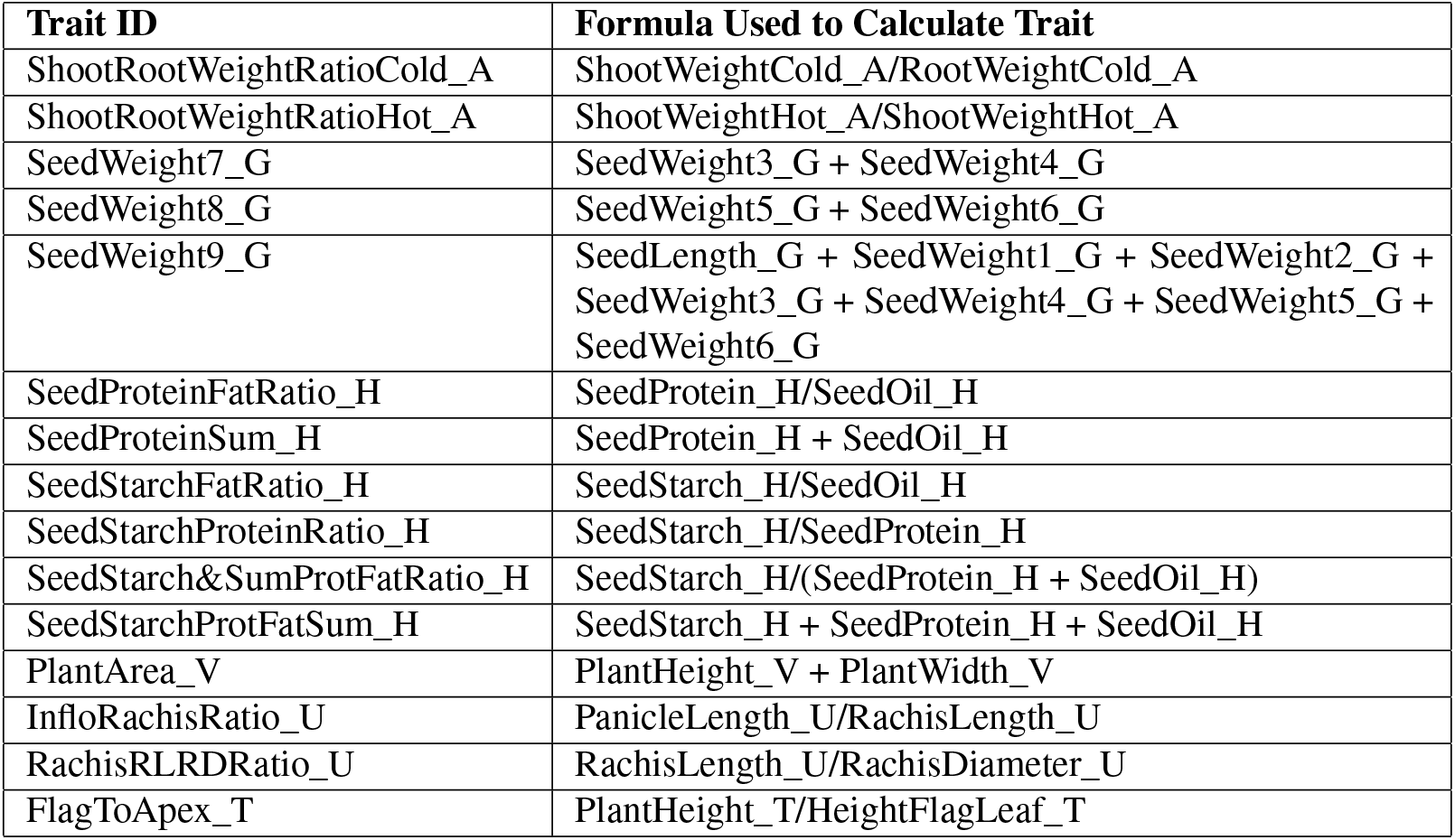
Formulas used to calculate derived traits.

| Trait ID | Formula Used to Calculate Trait |
| --- | --- |
| ShootRootWeightRatioCold_A | ShootWeightCold_A/RootWeightCold_A |
| ShootRootWeightRatioHot_A | ShootWeightHot_A/RootWeightHot_A |
| SeedWeight7_G | SeedWeight3_G + SeedWeight4_G |
| SeedWeight8_G | SeedWeight5_G + SeedWeight6_G |
| SeedWeight9_G | SeedLength_G + SeedWeight1_G + SeedWeight2_G + SeedWeight3_G + SeedWeight4_G + SeedWeight5_G + SeedWeight6_G |
| SeedProteinFatRatio_H | SeedProtein_H/SeedOil_H |
| SeedProteinSum_H | SeedProtein_H + SeedOil_H |
| SeedStarchFatRatio_H | SeedStarch_H/SeedOil_H |
| SeedStarchProteinRatio_H | SeedStarch_H/SeedProtein_H |
| SeedStarch&SumProtFatRatio_H | SeedStarch_H/(SeedProtein_H + SeedOil_H) |
| SeedStarchProtFatSum_H | SeedStarch_H + SeedProtein_H + SeedOil_H |
| PlantArea_V | PlantHeight_V + PlantWidth_V |
| InfloRachisRatio_U | PanicleLength_U/RachisLength_U |
| RachisRLRDRatio_U | RachisLength_U/RachisDiameter_U |
| FlagToApex_T | PlantHeight_T/HeightFlagLeaf_T |

